# Prefrontal cortex signals value category while basal ganglia represent learned values in value learning

**DOI:** 10.1101/2023.11.07.564561

**Authors:** M. Reza Heydari, Mohammad Ali Kheirkhah Ravandi, Okihide Hikosaka, Ali Ghazizadeh

## Abstract

Value learning can be variable for objects with the exact same reward but the underlying neural mechanism for such variability is not known. We have addressed this question by recording single-unit activity in the prefrontal cortex (PFC) and substantia nigra reticulata (SNr), two key nodes in cortex-basal ganglia circuitry with crucial roles in value learning, as macaque monkeys learned to associate novel objects with either high or low rewards. Estimating the trial by trial learned values based on choice performance, revealed stark differences between learned values across objects with the same reward outcome. Importantly, while PFC neurons rapidly learned to differentiate objects based on their value category, the firing in SNr correlated with the variability in learned values within a value category. Our results suggest that the variation in objects’ learned values is more likely to be a readout of SNr firing while PFC may provide a top-down teaching signal to basal ganglia to demarcate the value categories.

## Introduction

The ability to learn the values of novel objects in the environment is crucial for adaptability and ecological fitness. As the subject interacts with the objects and their reward outcomes, their learned values are gradually adjusted to reflect their true worth using the update rules in reinforcement learning formalism [1, 2, 3]. Nevertheless, it is often the case that different objects show different degrees of associability with reward (learned value) despite predicting the same explicit outcome (value category) even with similar training duration. This can be due to various internal factors such as fluctuating motivation levels or attention [4], different levels of uncertainty for the value of object [5] due to previous learning, generalization or interference in anterograde or retrograde directions [6, 7], or sensory salience of stimuli [8, 9]. Regardless of the cause, the neural mechanism that gives rise to the variability in learned values is not fully understood. It is also not known whether and how the veridical value category of objects can still be present in the brain despite the variation in learned values.

Neural correlates of value learning have been found in several brain structures in non-human primates, most notably in basal ganglia (BG) [10] and medial [11] as well as lateral prefrontal cortex (PFC) [12]. Caudate nucleus as an input nucleus of basal ganglia and in particular its head subregion (caudate head or CDh) is heavily involved in object value learning [13]. Caudate projects to substantia nigra pars reticulata (SNr) both directly and indirectly via external globus pallidus (GPe). SNr in turn projects to superior colliculus (SC), contributing to the control of saccadic eye movements guided by the value of objects in the visual scene [14]. Ventrolateral PFC (vlPFC) is also shown to rapidly learn and differentiate values of novel objects after a few repetitions [12]. Recent findings show that vlPFC and SNr which is the basal ganglia output have very similar coding capacity in signalling value memory during passive exposure to previously learned valuable objects [15]. However, it is not known whether this similarity extends to the active learning period and in particular whether the activity in these regions reflects trial by trial changes in learned values and their variability among objects.

To test this hypothesis, neural activity was recorded from vlPFC and SNr in macaque monkeys while they were exposed to novel fractal objects in each session and had to associate them with either high or low rewards. The subjects’ knowledge of object values was probed using sporadic choice trials which revealed faster learning for some objects compared to others in a session. Although value learning in vlPFC and SNr were initially similar, learning in vlPFC saturated early in a session, whereas the value signal in SNr continued to get larger and larger throughout the learning session. Notably, while SNr activity reflected the variability in learned values for the objects, vlPFC firing rapidly evolved to indicate the veridical value category of the objects regardless of learned value variation within a category.

## Results

Two monkeys (Monkey B and R) were trained to associate abstract visual objects (fractals) with either high or low rewards in a value learning task (Fig. 1A). Each value learning session was done with a novel set of eight fractals, four of which were randomly associated with high reward (good objects), and the other four were associated with low reward (bad objects). Neural activity during the task was recorded from vlPFC and SNr in separate sessions in each monkey. Learning sessions consisted of 160 trials (see materials and methods). In 80% of trials, a single good or bad object was presented peripherally and the monkey had to saccade to it to receive its associated reward (force trials). In 20% of trials, that were randomly spersed among forced trials, one good and one bad object were presented simultaneously and on diametrical positions on the screen (choice trials, see methods) (Fig. 1A). During the choice trials, the monkey had to choose one object by making a saccade to receive its reward. This resulted in on average 24 presentations for a given object across force and choice trials within a session.

**Figure 1:**
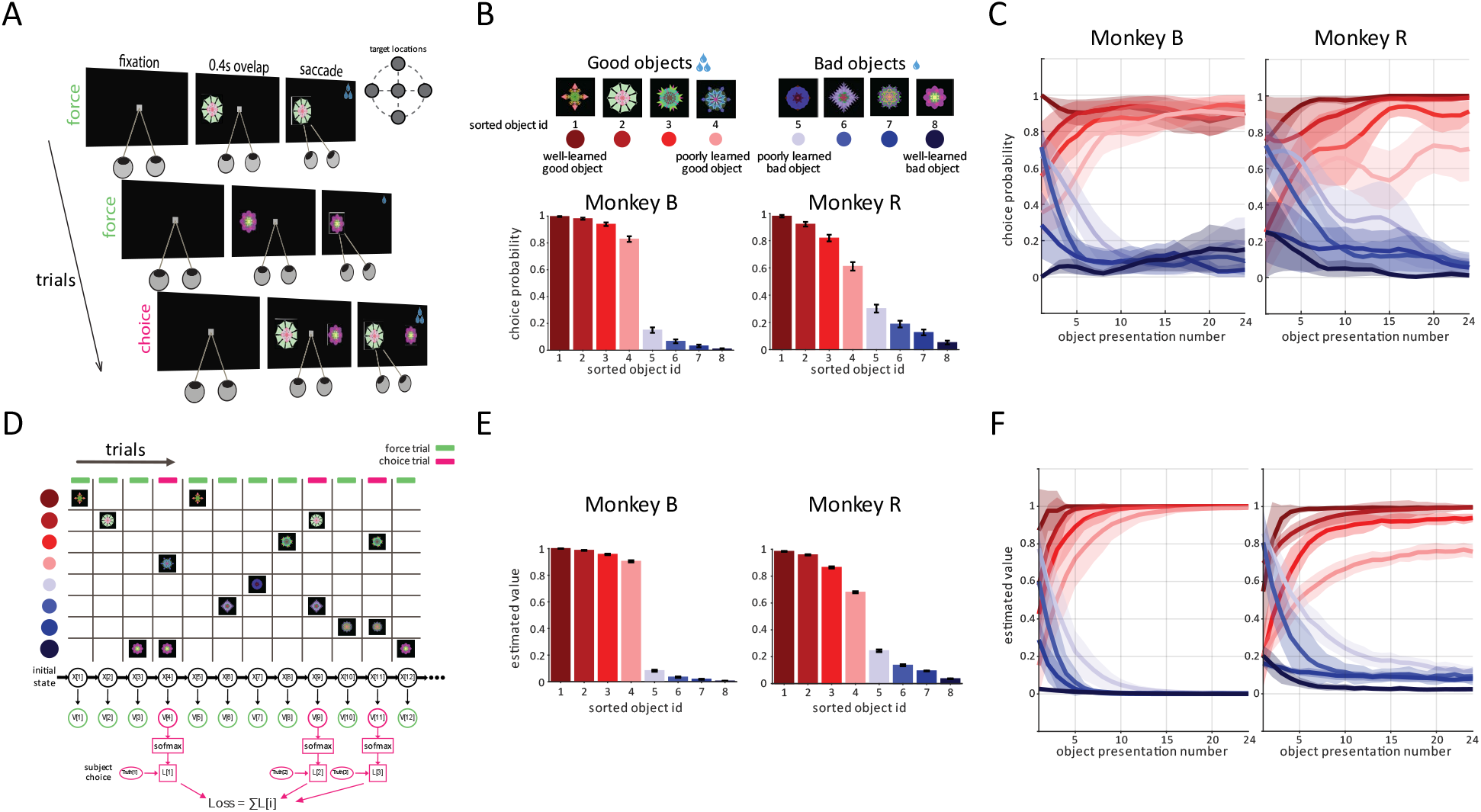
Different objects are learned with varying speeds as estimated by the RNN model. **(A)** Experimental paradigm: During each learning session, 8 novel fractals are presented in randomly across trials. Subjects are required to make a saccade to an object after fixation to receive the corresponding juice reward (high for good and low for bad) in force trials. In 20% of trials, the subject must choose between a good and a bad object. **(B)** Sorting objects by choice performance: The *8 objects within each session are sorted based on an overall measure of correct choices across all choice trials in the session (sorted object ID 1: the best learned good object. 8: the best learned bad object). (***C)** *The average trial by trial choice performance for sorted object IDs across sessions (note this trial by trial average is possible due to random appearance of choice trials during a session). The x-axis denotes the number of times an object is presented to the subjects from the start of the session. **(D)*** Values estimation with RNN model: *The RNN model calculates an estimated trial byt trial value for each object in each session using a Rescorla-Wagner like learning rule. Model parameters, such as initial value, learning and forgetting rates, are determined by minimizing the loss calculated from actual choice obserations and choice predictions from the model (see methods for details)* **(E)** Estimated values averaged for sorted object IDs across session. **(F)** *The estimated trial by trial values averaged for sorted objects and across all sessions. Errorbars and patches represent s.e.m here and throughout*.

Behavioral results show that both monkeys developed differential reaction times for saccading toward good compared to bad objects in force trials during training, consistent with previous findings [16, 12] (i.e. slower saccade to bad objects, Supp. Fig. 1A). Furthermore, both monkeys showed significant improvement in choice performance from chance level at 50% to >95% during a single learning session (Supp. Fig. 1B). The choice rate exhibited a nearly exponential learning curve and was faster in monkey B compared to monkey R (the trial constants for monkey B and R were 2.8 and 6.6, respectively).

Importantly, although all objects within a value category resulted in the same outcome (all good objects had the same high reward and all bad objects had the same low reward), results showed that the average choice performance for objects within good and bad object categories exhibited some variability (Fig. 1B, choice performance rank ordered and averaged across sets). The average choice performances across choice trials within a session revealed some well-learned and some poorly learned object values. In addition, the average trial-by-trial choice for objects (rank ordered by average choice in panel B) showed variability in the learning curves for individual objects (Fig. 1C). Note that due to random placement of choice trials within a session they could appear in any object presentation number allowing continuous curves for averages made across sessions.

While the trial by trial learned values of objects is not directly observed, one may estimate the learned values based on the sporadic choice performance within a session for each object (Fig. 1D) [17, 18, 19, 20]. We estimated the trial by trial subjective values that the monkey assigns to each object given the observed choice pattern and a simple value update model using a recurrent neural network (RNN) (see materials and methods). Consistent with the choice data the estimated learned values also showed variability between well-learned and poorly-learned good and bad objects across object sets (Fig. 1E-F).

Supp. Fig. 2 shows the firing activity of example neurons in vlPFC and SNr in monkeys B and R. Neurons in both regions developed differential responses to good and bad objects consistent with previous reports [21, 12, 22]. The SNr example neurons showed stronger excitation to bad compared to good objects while in vlPFC finding example neurons with higher firing to good (good-preferring, Gp) or higher firing to bad (bad-preferring, Bp) were equally likely. At the population level in SNr 2%, 81%, and 17% were Gp, Bp or showed no significant effect of value learning (non-significant or NS neurons) while in vlPFC 26%, 20%, and 54% were Gp, Bp or NS, respectively.

Average firing rate across neurons based on their value preference (see methods) on a trial by trial basis during learning showed that the differentiation between good and bad objects was predominantly due to a reduction of firing for the non-preferred value in vlPFC. In SNr, in addition to this response reduction to the non-preferred value, there was also some increase in response to the preferred value later in the session (Supp. Fig. 3). Across the population, the average value signal in each monkey rose rapidly in both vlPFC and SNr. After a few presentations, the value signal growth became slower in vlPFC while in SNr it kept rising until the end of learning. Notably, the value signal had a larger magnitude in monkey B compared to monkey R in particular in SNr. This neural difference was concurrent with faster value learning in monkey B as evident by choice rate and saccade reaction time as behavioral measures (Sup. Fig. 1, Fig. 1).

Notably and as can be seen in the example neurons in sup Fig.2, the neural responses in vlPFC were mostly the same for objects within a value category, while SNr firing showed a within category variability in firing rates. Population averages of neurons in each region based on the responses to preferred value further confirmed the striking difference in value coding scheme in vlPFC vs SNr (Fig. 2A). In vlPFC, there was a robust separation between preferred and non preferred value categories but there was little firing difference within a value category. On the other hand, in SNr, there was a graded response to objects based on their learned values with clear firing differences within a value category. We thus hypothesized that responses in vlPFC vs SNr encode value category vs learned value of objects, respectively (Fig. 2B). Consistently, the firing of SNr but not vlPFC neurons showed a significant positive correlation with objects’ learned values within both preferred and non-preferred value categories (Fig. 2C). In contrast, the neural responses in vlPFC while well separated for each value category did not positively correlate with learned values within a category. Indeed there was even a trending negative correlation in both preferred and non preferred categories (insets of Fig. 2C). This negative correlation was mostly caused by enhanced response difference to sorted object IDs 4 and 5 (poorly-learned objects in good and bad categories, respectively) which paradoxically had a larger difference compared to well-learned objects in each category (object IDs 1 and 8) possibly pointing to a prefrontal mechanism to accentuate the value category borders.

**Figure 2:**
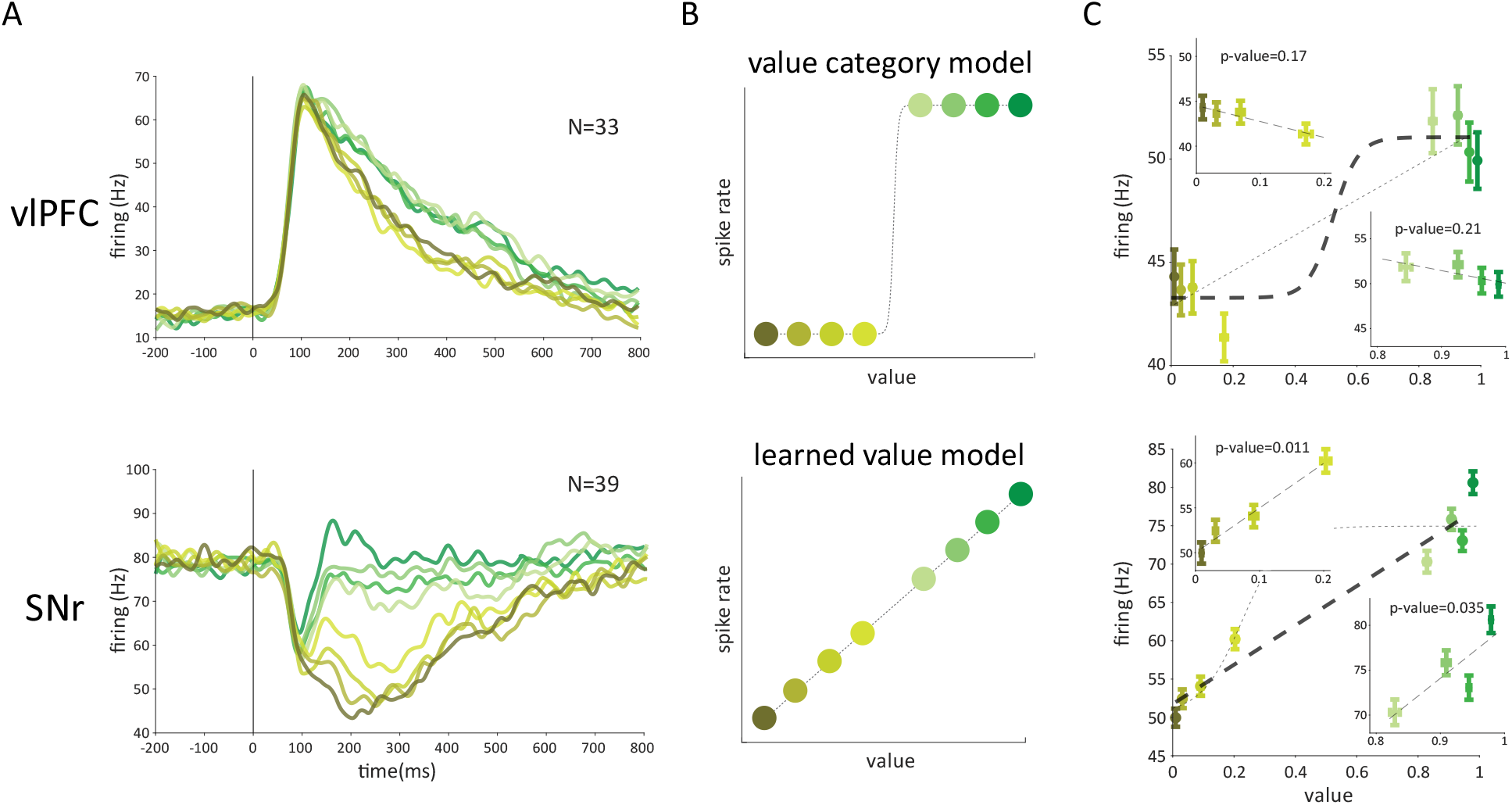
vlPFC represents value categories while SNr represents learned value during learning. **(A)** *The population firing rate averages for sorted object IDs based on neurons value preference time-locked to object onset, for vlPFC (top) and SNr (bottom). **(B)** Model of a hypothetical neuron encoding categorical value (top) vs learned-value (bottom) **(C)** Average neural response vs average estimated value for sorted object IDs. Two dashed lines denote fitted linear and sigmoid functions* (𝑅^2^ *for vlPFC: linear=0.8, sigmoid=0.85 and SNr: linear=0.93, sigmoid=0.84). Insets within the plots show the same information but separated for preferred and non-preferred objects (top-left: non-preferred, bottom-right: preferred). The significance of fitted linear slopes are noted*.

There could be several underlying reasons for the differential value learning within good and bad objects. One possibility is that differences in bottom-up physical features of the fractals may lead to better associability of some objects with high or low reward. If this was the case, one expects the sorted object ID for the same objects seen by both monkeys to be similar (i.e. a good object that is well-learned by Monkey B is also well-learned by Monkey R and vice versa). However, predicting the sorted ID of objects in one monkey based on the sorted ID of the same objects in the other monkey was not significantly different from the 25% chance level for 4 objects within each value category (Fig. 3A, 𝜒^2^test good: P=0.49 bad: P=0.58). Therefore, visual properties are not likely candidates for explaining the within-group differences in learned values between objects.

**Figure 3:**
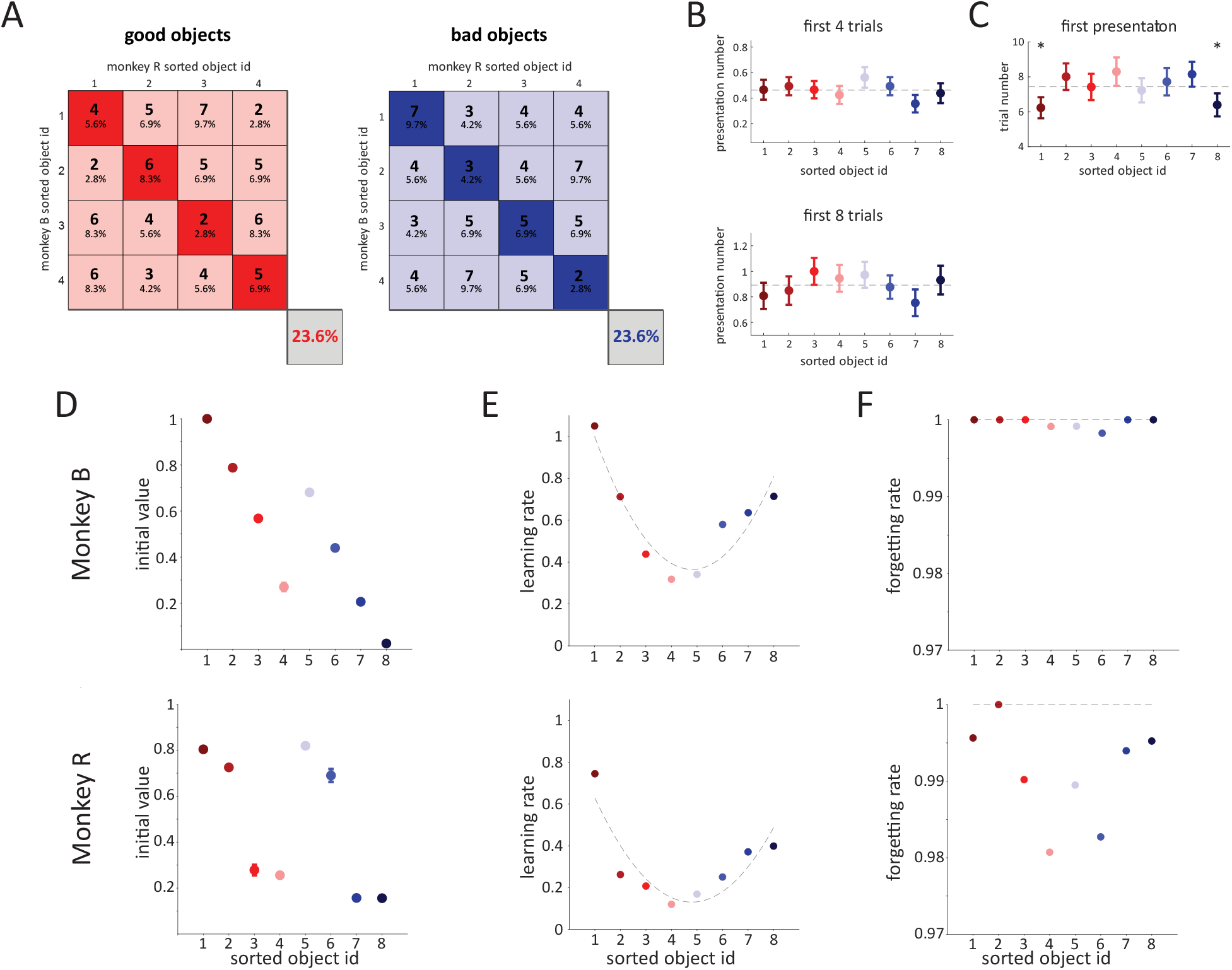
Factors that give rise to variability in learned value. **(A)** *The confusion matrix showing the consistency of sorted object ID for identical fractal objects between monkeys B and R. Each matrix entry shows the number and percentage (relative to all good or bad objects) of objects with a given sorted object IDs in Monkeys B vs R. Diagonal entries corresponds to objects with identical sorted object IDs for both monkeys. Off-diagonal entries refer to objects that have different sorted object IDs for monkeys. The bottom right percentages show the overall percentage of objects with the same sorted object ID between monkeys. **(B)** the number of times objects with distinct sorted object IDs were presented in the first 4 and 8 trials are depicted, Dashed lines are the means over the whole objects. **(C)** The first trial that on average a sorted object ID was seen in a session **(D)** The RNN parameter estimates of the initial value of sorted object IDs. **(E,F)** similar to D but for of learning rate (**E**) and forgetting rates (**F**) (dashed lines in **D** depict fitted parabola with* 𝑅^2^ *0.8 monkey B and 0.65 monkey R)*.

A second possibility is that the monkeys learned the value of objects that were presented more frequently in the first few trials of a session. However, looking at the encounter frequency did not reveal a systematic difference between objects in a category in the first 4 or 8 trials within a session (Fig. 3B, first 4 trials: F7,576=0.67 P=0.7, first 8 trials: F7,576=0.67 P=0.7). Nevertheless, we found that the best learned good and bad objects (sorted object IDs 1 and 8) had a tendency to be seen earlier than the other 6 objects within a session (Fig. 3C, F7,576=1.15 P=0.33, object 1 vs other 6: t113=2.3 P=0.022, object 8 vs other 6 t106=1.9 P=0.055).

A third possibility is that different objects may have started off the task with different initial values that could have facilitated or impeded their subsequent value learning depending on the consistency between these initial values and the outcomes presented in the task. Such deviations from neutral value could have been caused by value generalization mechanism from prior history with other fractals [5, 6, 7] or any other yet unspecified mechanism. This final possibility seems to be most consistent with our data as the initial estimated values for each object (Fig. 1F, the beginning of the presentation axis) showed variability that correlated well with the final learned values for that object in both monkeys (Pearson’s correlation, monkey B: good R=0.84 P=4e-45, bad R=0.78 P=1e-35, monkey R: good R=0.77 P=3e-25, bad R=0.78 P=4e-26). The average initial value of both within the good and bad object categories exhibited a consistent, near-monotonic decrease from well-learned objects to those that were poorly learned (Fig. 3D). In addition to the systematic difference in the initial values, we found a systematic difference in learning rates such that well-learned objects had much higher learning rates compared to poorly learned objects (Fig. 3E, F7,8=2.1 P=0.15). There was also a tendency for higher forgetting for poorly learned objects especially in monkey R (Fig. 3F, F7,8=0.48 P=0.83). Notably, using models that kept the initial value or learning/forgetting rates the same among objects, did not change our main finding about categorical value coding in vlPFC and learned value coding in SNr but models that allowed changes in initial values performed better in predicting choice performance in both monkeys based on BIC or AIC model selection criteria (Supp. Fig. 4, Supp. Table 1).

## Discussion

In many behavioral tasks, it is observed that the associability of objects with reward can show certain variations [4–9] (Fig. 1). It is not known which brain circuitry underlies this variability in learned value. We addressed this question using single unit recording in prefrontal cortex and basal ganglia output of nonhuman primates and by estimating the trial by trial learned values of objects. We found that SNr firing which represents the basal ganglia output reflected variabilitity in learned values for objects. On the other hand, vlPFC neurons did not encode this variation in learned values and instead rapidly learned to signal the value category of objects based on the received outcome (Fig. 2). Therefore the variation in objects’ learned values which was manifest in choice behavior is more likely to be a result of variability in SNr firing rather than vlPFC activity.

Both SNr and vlPFC have been previously shown to encode the value of objects quite similarly with respect to the reward granularity and uncertainty [15]. That is when the actual outcomes themselves show variation, vlPFC tracks those external variations similarly to SNr. This effect is consistent with the current findings as graded reward created multiple value categories all of which are tracked by vlPFC. Nevertheless, it was also shown that during a passive viewing task, where subjects were required to maintain their gaze on the central fixation point while objects were displayed instead of making saccades toward objects, variations in value memory of objects with the same outcome were encoded in a correlated fashion in SNr and vlPFC [15]. This latter fact may seem at odds with our current finding that the variation in learned value during the training was correlated with firing in SNr but not with firing in vlPFC. One possibility is that the neurons in vlPFC can signal value category and ignore variation in learned value during the training task but then switch to reflect the variation of value memory during the passive viewing task. Indeed there is evidence that shows that unlike SNr the value preference of vlPFC neurons can switch between value learning and passive viewing tasks (Supp. Fig. 5). This could also be due to the fact that value training was an active task which required a saccade while in the passive task monkey had to inhibit saccades to maintatin central fixation. Nevertheless the curious difference between the roles of vlPFC in active and passive tasks requires further investigation.

Our findings are in agreement with previous research implicating the role of vlPFC in categorizing objects [23], and the role of SNr in habit and object skill learning [24, 16, 25]. In such a scheme, one can imagine that in our value learning task, vlPFC rapidly creates a top-down demarcation of good and bad object categories. On the other hand, habitual learning happens in a more gradual, step-by-step fashion that continues for quite a long time. In this case, SNr is involved in more automatic and habitual attraction towards good objects. Indeed, our results show that vlPFC rises sharply in the first few trials and saturates soon. In contrast, the value signal in SNr continues to rise monotonically, without being saturated after tens of trials (Supp. Fig. 3.).

vlPFC and SNr are reciprocally connected to each other in the cortico-basal ganglia-thalamo-cortical loop [26]. Based on our results, one possible function of this loop could be for vlPFC to provide a veridical teaching signal about value categories to SNr. Consistent with this hypothesis, we find that vlPFC firing may work to accentuate the category differences for poorly learned objects (Fig. 2C). Furthermore, our results show that the variations in learned values stem from nonequal initial values and learning rates for novel objects. SNr firing could then be influenced by these initial differences in object values as well as the value categories that are signaled by vlPFC. Direct causal maniplutions of activities in vlPFC and SNr can further delinate their role in value learning and their impact on each other in the future.

## Materials and methods

### Subjects and surgery

Two male adult rhesus monkeys were used in all tasks (monkeys B and R ages 7 and 10, respectively). All animal care and experimental procedures were approved by the National Eye Institute Animal Care and Use Committee and complied with the Public Health Service Policy on the humane care and use of laboratory animals. Both animals underwent surgery under general anesthesia during which a head holder and recording chambers were implanted on the head and scleral search coils for eye tracking were inserted in the eyes. After confirming the position of recording chamber using MRI, craniotomies over PFC and cdlSNr on the same hemisphere were performed during a second surgery. For detail description refer to [15].

### Recording localization

Substantia nigra (SN) recording localization in both subjects was done using T1- and T2-weighted MRI (4.7 T, Bruker). T2-weighted MRI is especially useful for imaging SNr area due to higher iron content [27]. During imaging, the recoding chambers were equipped with a grid with 2 mm hole spacing and were filled with gadolinium for better contrast. The location of SN in each monkey was further verified using the standard monkey atlas (D99 atlas [28]) which was brought into each monkeys native space using the NMT toolbox [29] and the projection of SN as reachable through the posterior recording chamber was visualized and confirmed to coincide with the recording locations (refer to Supp. Fig. 1 in [15]). vlPFC recordings were localized and reconstructed using the same scans per monkey and using the D99 atlas (refer to Supp. Fig. 1 in [12]).

### Stimuli

Visual stimuli with fractal geometry were used as objects [30]. One fractal was composed of four point-symmetrical polygons that were overlaid around a common center such that smaller polygons were positioned more toward the front. The parameters that determined each polygon (size, edges, color, etc.) were chosen randomly. Fractal diameters were on average ∼7° (ranging from 5°−10°). Monkey B saw 576 novel objects (344 objects of which was along with recording) and monkey R saw 456 novel objects (248 along with recording) in value training task.

### Task control and neural recording

All behavioral tasks and recordings were controlled by a custom written VC++ based software (Blip; http://www.robilis.com/blip/). Data acquisition and output control was performed using National Instruments NI-PCIe 6353. Eye position was sampled at 1 kHz using scleral search coils. Diluted apple juice (33% and 66% for monkey B and R respectively) was used as a reward. Activity of single isolated neurons was recorded with acute penetrations of glass coated tungsten electrodes (Alpha-Omega, 250μm total thickness). The electric signal from the electrode was amplified and filtered (2 Hz-10 kHz) and was digitized at 1 kHz. Neural spikes were isolated online using voltage-time discrimination windows. The results reported are from a total of 33 vlPFC neurons(23 and 10 in monkeys B and R) and from 39 SNr neurons (18 and 21 in monkeys B and R). For more recording details refer to [15].

### Value training task

Each session of training was performed with one set of fractals, which was not seen before. The good/bad sets consisted of 8 fractals (4 good / 4 bad fractals). Bad fractals were paired with low reward (0.07 ml and 0.1 ml in monkeys R and B, respectively) and good fractals were paired with high reward (0.2 ml and 0.3 ml for monkeys R and B, respectively). The different juice amount was customized for each monkey based on his water motivation and to ensure satisfactory cooperation. The high to low juice amount was about 3 to 1 in both subjects. In a given trial, after central fixation on a white dot (2°) one object appeared on the screen at one of the five peripheral locations (10–15° eccentricity) or at the center. After an overlap of 400ms, the fixation dot disappeared and the animal was required to make a saccade to the fractal. After 500 ± 100ms of fixating the fractal, a large or small reward was delivered. The displayed fractal was then turned off followed by an inter-trial intervals (ITI) of 1–1.5 s with a black screen. Breaking fixation or a premature saccade to fractal during the overlap period resulted in an error tone (monkey B: 13% and 18%, monkey R: 13% and 27% of force and choice trials). A correct tone was played after a correct trial. Normally a training session with neural recording consisted of 160 trials (range spanning from 60 to 173 trials) with objects presented in random order. For sessions focusing solely on behavior measurements, there were 80 trials. To check the behavioral learning of object values, choice trials with two objects with different reward association were included randomly in one out of five trials (20% of trials). During the choice the two fractals were shown in diametrically opposite locations and monkey was required to choose one by looking and holding gaze for 500 ± 100 ms on a fractal after which both fractals were turned off and the corresponding reward would be delivered. Only a single saccade was allowed in choice trials. The location and identity of fractals were randomized across choice trials. Throughout this document, the term “presentation number” refers to the instances from the beginning of a session in which each object was observed, including both force and choice correct trials.

### Passive viewing task

A passive viewing trial started after central fixation on a white dot (2°). The animal was required to hold a central fixation while objects from a given set were displayed randomly with 400 ms on and 400 ms off schedule. Animal was rewarded for continued fixation after a random number of 2–4 objects were shown. A trial would abort if animal broke fixation or made a saccade to objects (<1% of trials). Objects were shown close to the location with maximal visual response for each neuron as determined by receptive field mapping task. When this maximal location was close to center (<5°) passive viewing was sometimes done by showing objects at the center. A block of passive viewing consisted of 5-6 presentations per object. In most cases,more than one block was acquired for a given set. For detail description and analysis of this task refer to [15].

### Data analysis

Neural firing rates were filtered using MATLAB *filtfilt* with a 10 ms bandwidth after spike counting within a 20 ms time window. Responses were time-locked to object-onset for analysis in all tasks. The analysis epoch was spanned from 200 ms to 600 ms after object onset for calculation of AUC and assign a single neural response per trial. The good-bad discrimination of each neuron was quantified by measuring the average firing during analysis epoch across trials using AUC, and significance levels were determined using the Wilcoxon rank-sum test. MATLAB *glmfit* with default parameters were used to fit linear models and significance evaluation (Fig. 2C, Supp. Fig. 4B). For sigmoid (Fig. 2C, Supp. Fig. 4B), exponential (Supp. Fig. 1B), and parabola (Fig. 3D) curve fittings, custom functions with an appropriate parameter sets were defined, and parameters were tuned by minimizing MSE loss between predictions and responses.

PYTHON *tensorflow* were used to define the RNN network with the customized architecture. The model was trained using the ADAM optimizer, a learning rate of 0.01, and 2000 epochs. A train-test split of 80%-20% was applied, and no case of overfitting was observed (maximum ∼8% loss difference measured). The ANOVA test for the signifance evaluation of the learning/forgetting rates (Fig. 3D-E) was calculated by aggregating values across both monkeys B and R.

### RNN for value estimation

In the RNN model, the value of each fractal is represented by a time-dependent process *V_i_^s^*[*k*], in which 𝑖 ∈ {1, …, 8} is the index of the sorted object ID of good and bad object, 𝑠 is the session number, and 𝑘 ∈ {1, …, 160} corresponds to each trial in session 𝑠. These values can be viewed as the output variable of a RNN network, which results from a projection of the hidden state variables *X_i_^s^*[*k*]. Each state variable *X_i_^s^*[*k* + 1] is influenced by the preceding time step variable *X_i_^s^*[*k*] through an unknown underlying process. The goal of the RNN model is to discover this dynamic process and aforementioned projection by the aim of the choice of the monkey in choice trials.

There are two potential scenarios for the 𝑖th object in the 𝑘th trial of session 𝑠. First scenerio, the object is viewed either in a choice or force trial. In this case, the value of the object undergoes an update based on Rescorla-Wagner’s delta rule [3]. The value moves towards its final value by incorporating a positive learning rate and the current error between the final value and the current value. Second scenario is when the object is seen neither in a choice nor force trial, in which case its value decays towards zero by being multiplied by a forgetting rate. Putting them together, the dynamical process can be described as follows:

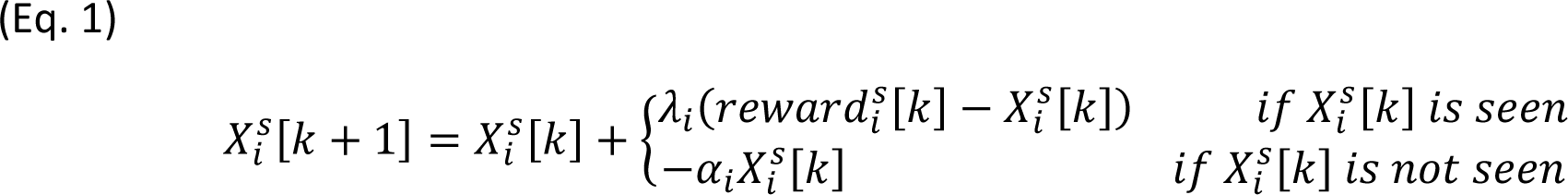

*reward^s^_i_* is the reward associated with the observed object, taking the value of -1 for bad objects and 1 for good objects. 𝜆_𝑖_ and 1 − 𝛼_𝑖_are learning and forgetting rates, respectively. Note that we used a different learning rate parameter for objects sorted by their average choice performance during a session. We refer to this as sorted object ID going from 1-4 for best to worst learned good object and from 5-8 for worst to best learned bad object (Fig. 1B). For instance, we used the same learning and forgetting rate for all objects with sorted ID 1 across all sessions for each monkey. The same holds for different initial values 𝑋_𝑖_[0] for each of objects. In each trial 𝑘, only one object is observed and the other objects are forgotten (Unless for choice trials where a good and bad object is updated by the first line of Eq. 1).

The output projection for the value calculation is based on the Eq. 2. This relationship suppress the input values so that the estimated value is between 0: lowest value and 1: highest value.

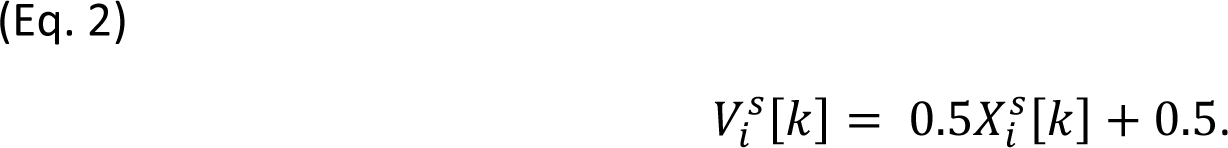

Two constraints 𝜆_𝑖_ > 0 and 0 ≤ 𝛼_𝑖_ ≤ 1 ensure that the values 𝑉^𝑠^[𝑘] remain within the range [0, +1]. The complete model consists of only three sets of parameters 𝜆_𝑖_, 𝛼_𝑖_, and the initial value 𝑋_𝑖_[0] (in total 24 parameters per monkey).

To train the model, choice trials were used as the ground truth for values of the two seen good and bad objects. We computed an estimated choice probability of both 𝑚th good and 𝑠th bad objects with a Softmax function on their estimated values at choice trial 𝑘 in session 𝑠 (Eq. 3)

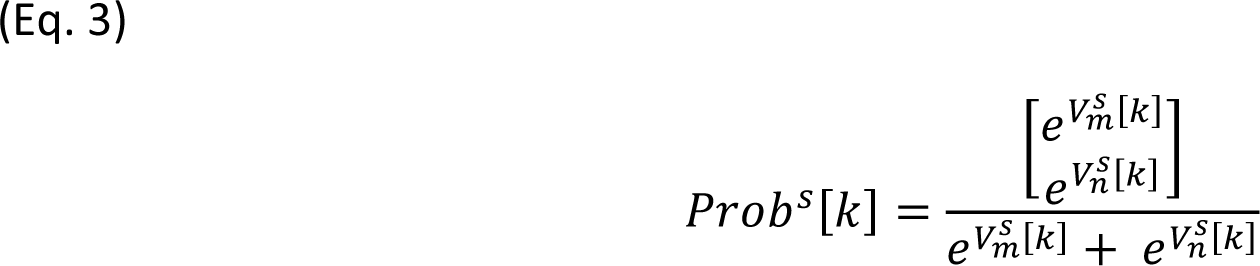

This vector represents how much it is possible for the subject to choose between two objects 𝑚 and 𝑠. The cross entropy loss was then calculated between this estimated choice probability and the real choice of the monkey (Eq. 4) for session 𝑠 at trial 𝑘.

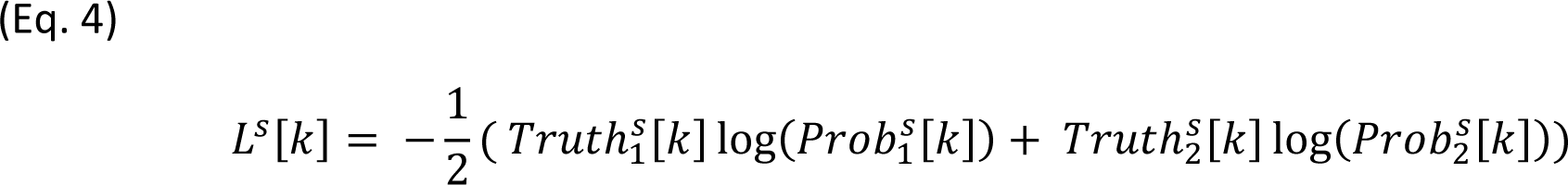

Total loss is the above loss averaged over all sessions (𝑆: number of sessions): (Eq. 5)

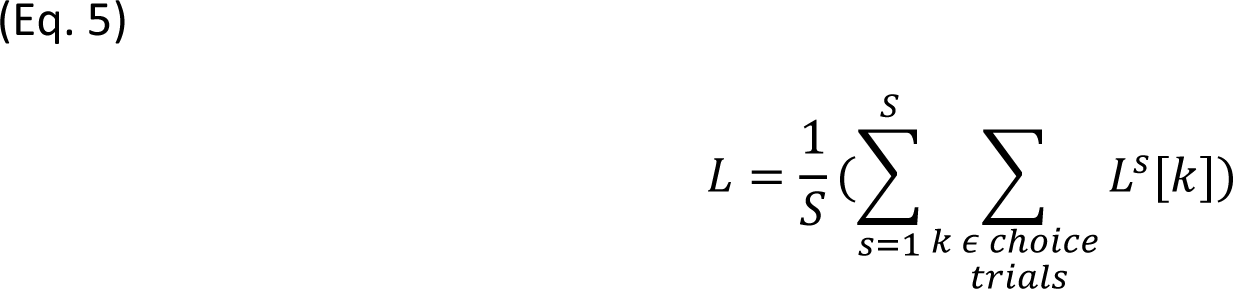

By the end of the training procedure, estimated values 𝑉^𝑠^[𝑘] were obtained for each sorted object ID at each trial of a given session.

Other scenerios where only the initial values but not the learning/forgetting factors were different among the object sorted IDs or conversely only the leanring/forgetting factors but not the initial values were different among the sorted object IDs were also tested (Supp. Fig. 4A). Our main conclusions about the role of vlPFC and SNr did not depend on the choice of model (Supp. Fig. 4B). The best-performing model was determined based on AIC/BIC scores (Supp. Table 1). For the sake of consistency model 1 is used for both monkeys for the main results presented throughout the paper. The parameter estimation for each monkey was done using all behavioral session regardless of whether recording was done (129 sessions in total, 74 sessions with recording).

### Statistical tests and significance levels

One-way ANOVA were used to test main effects. Note that since there is a single learning/forgetting rate for each sorted object ID in each monkey (Supp. Fig. 3E, F), the values are combined across both monkeys to compute the ANOVA test. Error-bars in all plots show standard error of the mean (s.e.m). Significance threshold for all tests in this study was 𝑝 < 0.05.

---

**Supplementary Figure 1:**
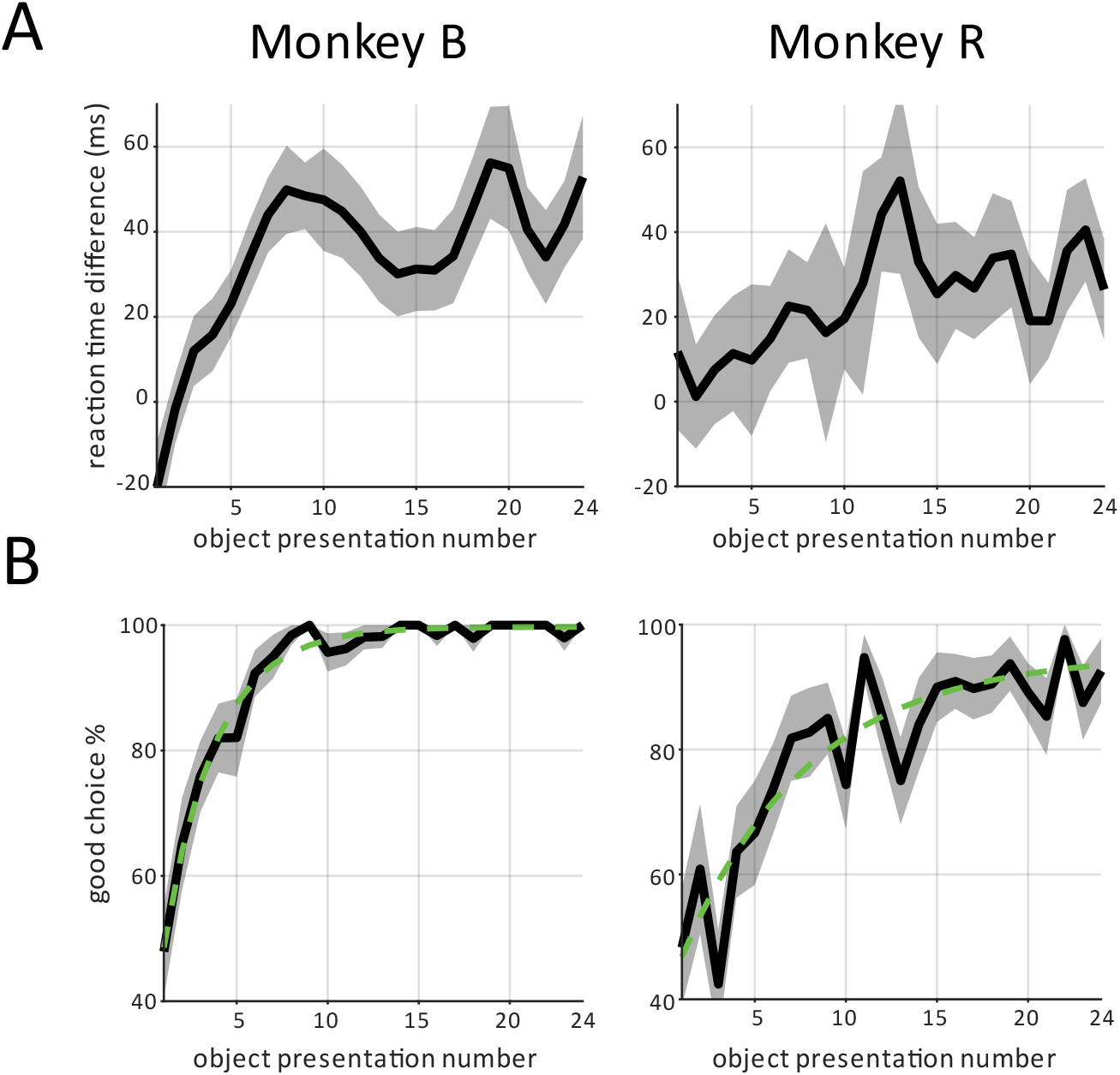
The value learning and behavioral measures. **(A)** The difference in reaction time between bad and good fractals during force trials. **(B)** Subjects learn to choose good objects from ∼50% level at the start of fractal presentation to 99.7% (B) and 94.3% (R). Green dashed lines depict fitted exponential curves (𝑅^2^ is 0.98 for B and 0.83 for R).

**Supplementary Figure 2:**
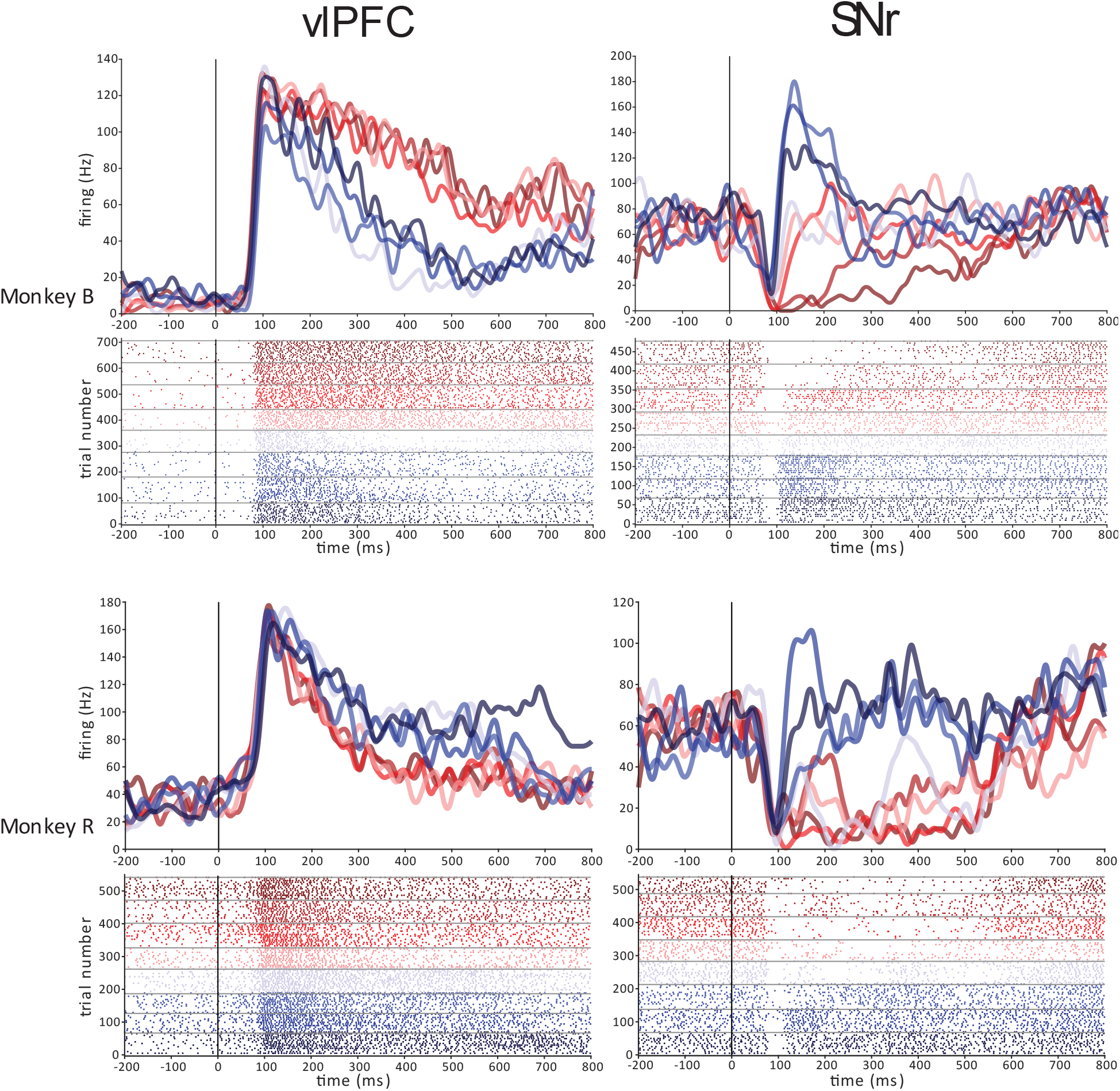
Value coding in examples neurons. PSTH and raster plots of one example neuron from each region (left: vlPFC, right: SNr) in each monkey (top: B, bottom: R). Both SNr neurons are bad-preferring (Bp). vlPFC example on top is good-preferring (Gp), while the bottom vlPFC neuron is Bp. Sorted object IDs are separated by different color codes as is in Figure 1B.

**Supplementary Figure 3:**
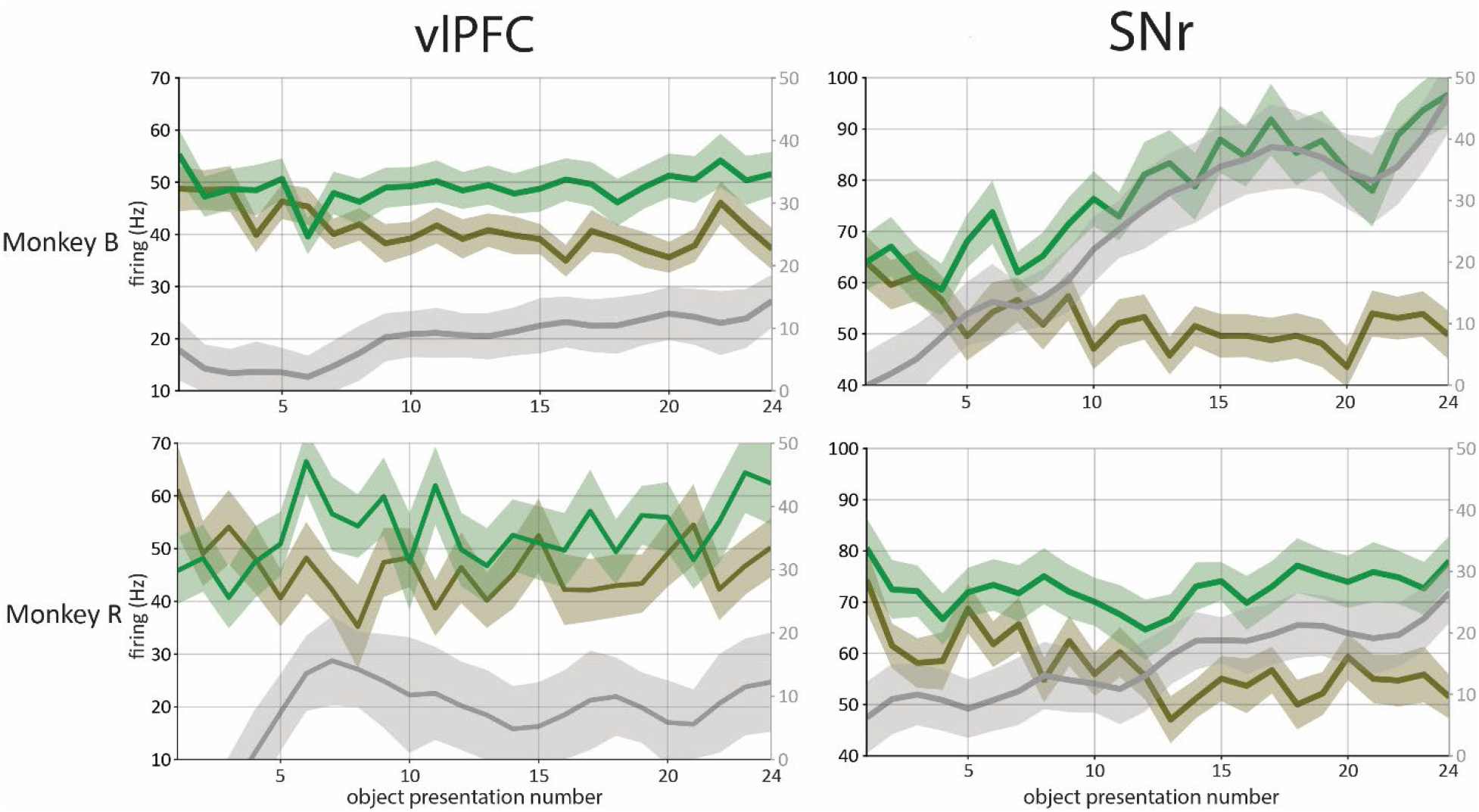
Trial by trial value learning in vlPFC and SNr. Average neural firing activity is displayed for each region. Neuronal responses are categorized based on the value preference of each neuron: preferred (green, representing good responses of Gp neurons and bad responses of Bp neurons) and non-preferred (brown, indicating bad responses of Gp neurons and good responses of Bp neurons) values in force trials. The grey traces on the right y-axis show the smoothed difference (using filtfilt with window size of 3) between the responses to preferred and non-preferred stimuli. The x-axis denotes the number of times an object is presented to the subjects from the start of the session.

**Supplementary Figure 4:**
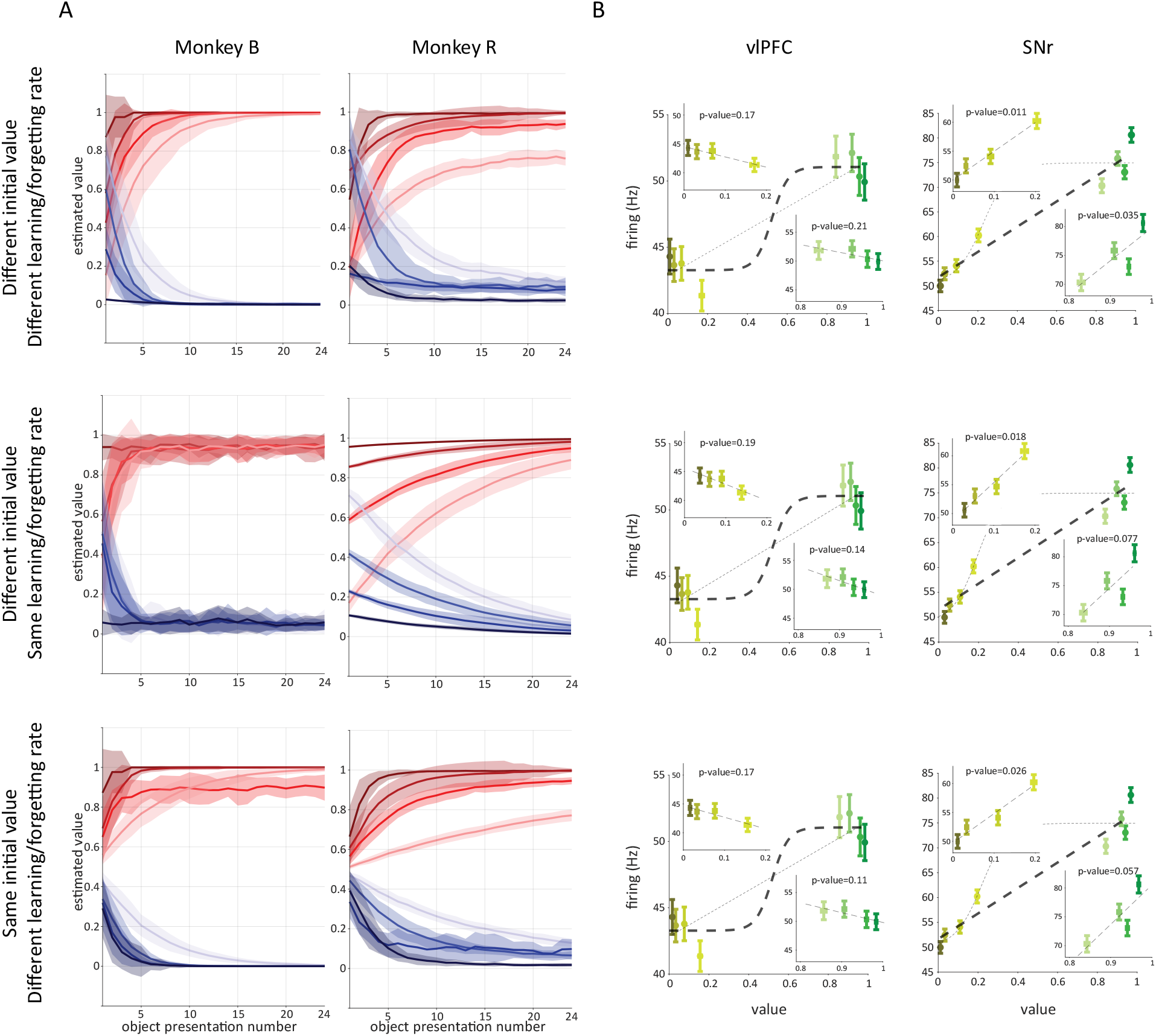
Comparison of different RNN models. We trained RNN model with three distinct set of parameters: Different initial values for sorted object IDs with different learning and different forgetting rates (top row, the same as the model used in the main paper). Different initial values for sorted object IDs but the same learning and forgetting rates for all objects (middle row). The same initial value but different learning and different forgetting rates for each sorted object IDs (bottom row). **A** and **B** are the same as the figures 1F and 2B.

**Supplementary Table 1:** Comparison of different RNN models based on AIC/BIC

| Model | Monkey B |  | Monkey R |  |
| --- | --- | --- | --- | --- |
|  | AIC | BIC | AIC | BIC |
| Different initial value<br>Different learning/forgetting rate | **525.4** | **692.7** | 995.9 | 1088.6 |
| Different initial value<br>Same learning/forgetting rate | 626.7 | 725 | **981.4** | **1076** |
| Same initial value<br>Different learning/forgetting rate | 605.4 | 700 | 1004.5 | 1164 |

**Supplementary Figure 5:**
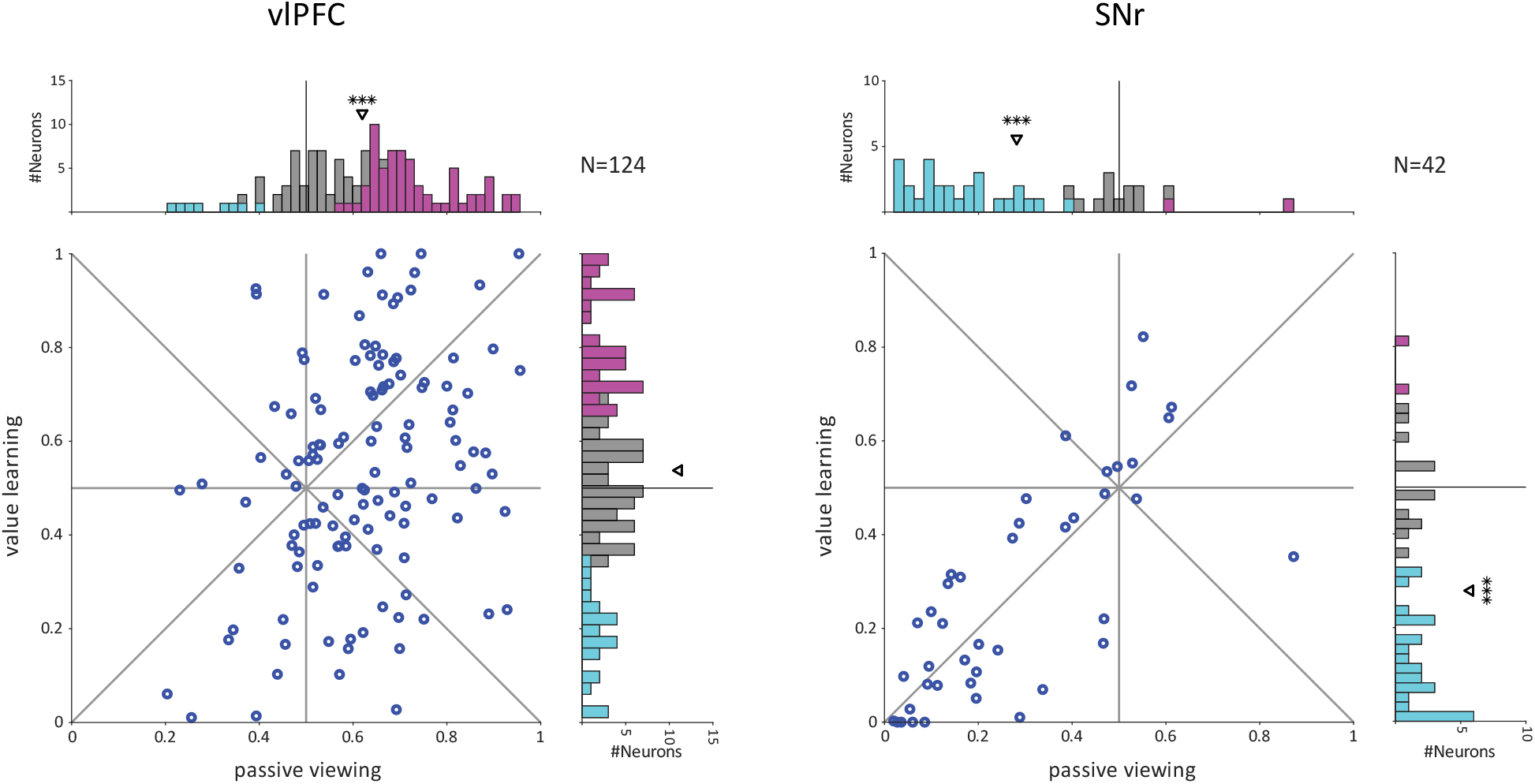
Neural object representation in vlPFC and SNr during value learning requiring saccades and passive viewing task. Each dot in scatter plots represents AUC of good-bad differentiation among vlPFC (left, 124 units) and SNr (right, 42 units) neurons for both value learning (y-axis) and passive viewing (x-axis) tasks. The distribution of good vs bad discrimination is depicted on the right (value learning, vlPFC: mean=0.54 P=0.078, SNr: mean=0.28 P=2e-7) and on the top (passive viewing, vlPFC: mean=0.62 P=5e-14, SNr: mean=0.28 P=2e-8) of each scatter plot. Results show consistent value coding in SNr but not in vlPFC.

## References

[1] R. S. Sutton and A. G. Barto, Reinforcement Learning: An Introduction, 1998.

[2] R. S. Sutton and A. G. Barto, “Toward a modern theory of adaptive networks: expectation and prediction.,” Psychological review, vol. 88, no. 2, p. 135, 1981.

[3] R. A. Rescorla and A. R. Wagner, “A theory of Pavlovian conditioning: The effectiveness of reinforcement and non-reinforcement.,” AH Black & WF Prokasy,, pp. 64–69, 1972.

[4] Mackintosh and N. J, “A theory of attention: Variations in the associability of stimuli with reinforcement.,” Psychological review, vol. 82, no. 4, p. 276, 1975.

[5] P. Dayan and A. J. Yu, “Uncertainty and learning.,” IETE Journal of Research, vol. 49, no. 2.3, pp. 171-181, 2003.

[6] S. Kakade and P. Dayan, “Dopamine: generalization and bonuses.,” Neural Networks, vol. 15, no. 4-6, pp. 549–559, 2002.

[7] W. Schultz, “Predictive reward signal of dopamine neurons.,” Journal of neurophysiology, 1998.

[8] I. P. Pavlov, “Conditioned Reflexes,” *London*: Oxford University Press, 1927.

[9] D. D. Foree and V. M. LoLordo, “Attention in the pigeon: differential effects of food-getting versus shock-avoidance procedures.,” Journal of Comparative and Physiological Psychology, vol. 85, no. 3, p. 551, 1973.

[10] O. Hikosaka, H. F. Kim, M. Yasuda and S. Yamamoto, “Basal ganglia circuits for reward value-guided behavior,” Annu Rev Neurosci, vol. 37, pp. 289–306, 2014.

[11] P. H. Rudebeck and E. L. Rich, “Orbitofrontal cortex.,” Current Biology, vol. 28, no. 18, pp. R1083-R1088, 2018.

[12] A. Ghazizadeh, S. Hong and O. Hikosaka, “Prefrontal cortex represents long-term memory of object values for months,” Current Biology, vol. 28, no. 14, pp. 2206-2217. e5, 2018.

[13] O. H. Hyoung F. Kim, “Distinct Basal Ganglia Circuits Controlling Behaviors Guided by Flexible and Stable Values,” Neuron, vol. 79, no. 5, pp. 1001-1010, 2013.

[14] a. R. H. W. O. Hikosaka, “Visual and oculomotor functions of monkey substantia nigra pars reticulata. I. Relation of visual and auditory responses to saccades,” Journal of Neurophysiology, vol. 49, no. 5, 1983.

[15] A. Ghazizadeh and O. Hikosaka, “Common coding of expected value and value uncertainty memories in the prefrontal cortex and basal ganglia output,” Science Advances, vol. 7, no. 20, p. eabe0693, 2021.

[16] H. F. Kim, A. Ghazizadeh and O. Hikosaka, “Dopamine neurons encoding long-term memory of object value for habitual behavior,” Cell, vol. 163, no. 5, pp. 1165-1175, 2015.

[17] M. Usher and J. L. McClelland, “The time course of perceptual choice: the leaky, competing accumulator model.,” Psychological review, vol. 108, no. 3, p. 550, 2001.

[18] W. Schultz, “Neuronal reward and decision signals: from theories to data.,” Physiological reviews, vol. 95, no. 3, pp. 853–951, 2015.

[19] B. M. Turner, B. U. Forstmann, E. J. Wagenmakers, S. D. Brown, P. B. Sederberg and M. Steyvers, “A Bayesian framework for simultaneously modeling neural and behavioral data.,” NeuroImage, vol. 72, pp. 193–206, 2013.

[20] G. de Hollander, B. U. Forstmann and S. D. Brown, “Different ways of linking behavioral and neural data via computational cognitive models.,” Biological Psychiatry: Cognitive Neuroscience and Neuroimaging, vol. 1, no. 2, pp. 101–109, 2016.

[21] M. Yasuda, S. Yamamoto and O. Hikosaka, “Robust representation of stable object values in the oculomotor basal ganglia.,” Journal of Neuroscience, vol. 32, no. 47, pp. 16917-16932, 2012.

[22] O. Hikosaka, H. F. Kim, H. Amita, M. Yasuda, M. Isoda, Y. Tachibana and A. Yoshida, “Direct and indirect pathways for choosing objects and actions.,” European Journal of Neuroscience, vol. 49, no. 5, pp. 637–645, 2019.

[23] M. R. T. P. a. E. K. M. David J. Freedman, “Visual Categorization and the Primate Prefrontal Cortex: Neurophysiology and Behavior,” Journal of Neurophysiology, vol. 88, no. 2, 2002.

[24] M. Isoda and O. Hikosaka, “Cortico-basal ganglia mechanisms for overcoming innate, habitual and motivational behaviors.,” European Journal of Neuroscience, vol. 33, no. 11, pp. 2058-2069, 2011.

[25] O. Hikosaka, S. Yamamoto, M. Yasuda and H. F. Kim, “Why skill matters.,” Trends in cognitive sciences, vol. 17, no. 9, pp. 434–441, 2013.

[26] G. E. Alexander, M. R. DeLong and P. L. Strick, “Parallel organization of functionally segregated circuits linking basal ganglia and cortex.,” Annual review of neuroscience, vol. 9, no. 1, pp. 357–381, 1986.

[27] T. Hashido and S. Saito, “Quantitative T1, T2, and T2* mapping and semi-quantitative neuromelanin-sensitive magnetic resonance imaging of the human midbrain.,” PloS one, vol. 11, no. 10, p. e0165160, 2016.

[28] C. Reveley, A. Gruslys, F. Q. Ye, D. Glen, J. Samaha, B. E. Russ, Z. Saad, A. K. Seth, D. A. Leopold and K. S. Saleem, “Three-dimensional digital template atlas of the macaque brain.,” Cerebral cortex, vol. 27, no. 9, pp. 4463-4477, 2017.

[29] J. Seidlitz, C. Sponheim, D. Glen, Q. Y. Frank, K. S. Saleem, D. A. Leopold, L. Ungerleider and A. Messinger, “A population MRI brain template and analysis tools for the macaque.,” Neuroimage, vol. 170, pp. 121–131, 2018.

[30] Y. Miyashita, S.-I. Higuchi, K. Sakai and N. Masui, “Generation of fractal patterns for probing the visual memory.,” Neuroscience research, vol. 12, no. 1, pp. 307–311, 1991.

[31] “Dopaminergic Neurons and Brain Reward Pathways,” The American Journal of Pathology, vol. 186, no. 3, p. 478–488, 2016.

[32] “Functional Neuroanatomy of the Basal Ganglia,” Cold Spring Harbor Perspectives in Medicine, vol. 2, no. 12, 2012.

[33] “Glutamate as a neurotransmitter in the healthy brain,” Journal of Neural Transmisson, vol. 121, no. 8, pp. 799–817, 2014.

[34] O. Hikosaka, Y. Takikawa and R. Kawagoe, “Role of the basal ganglia in the control of purposive saccadic eye movements,” Physiol Rev, vol. 800, no. 3, pp. 953–78, 2000.

[35] O. Hikosaka and R. H. Wurtz, “Visual and oculomotor functions of monkey substantia nigra pars reticulata. I. Relation of visual and auditory responses to saccades,” J Neurophysiol, vol. 49, no. 5, pp. 1250-53, 1983.

[36] T. K. Moon, “The expectation-maximization algorithm,” IEEE Signal Processing Magazine, vol. 13, no. 6, pp. 47–60, 1996.

[37] O. Hikosaka, K. Nakamura and H. Nakahara, “Basal ganglia orient eyes to reward.,” Journal of neurophysiology, vol. 95, no. 2, pp. 567–584, 2006.

