## Supplementary figures and images for "Prefrontal cortex signals value category while basal ganglia represent learned values in value learning"

### Supplemental Figure 1

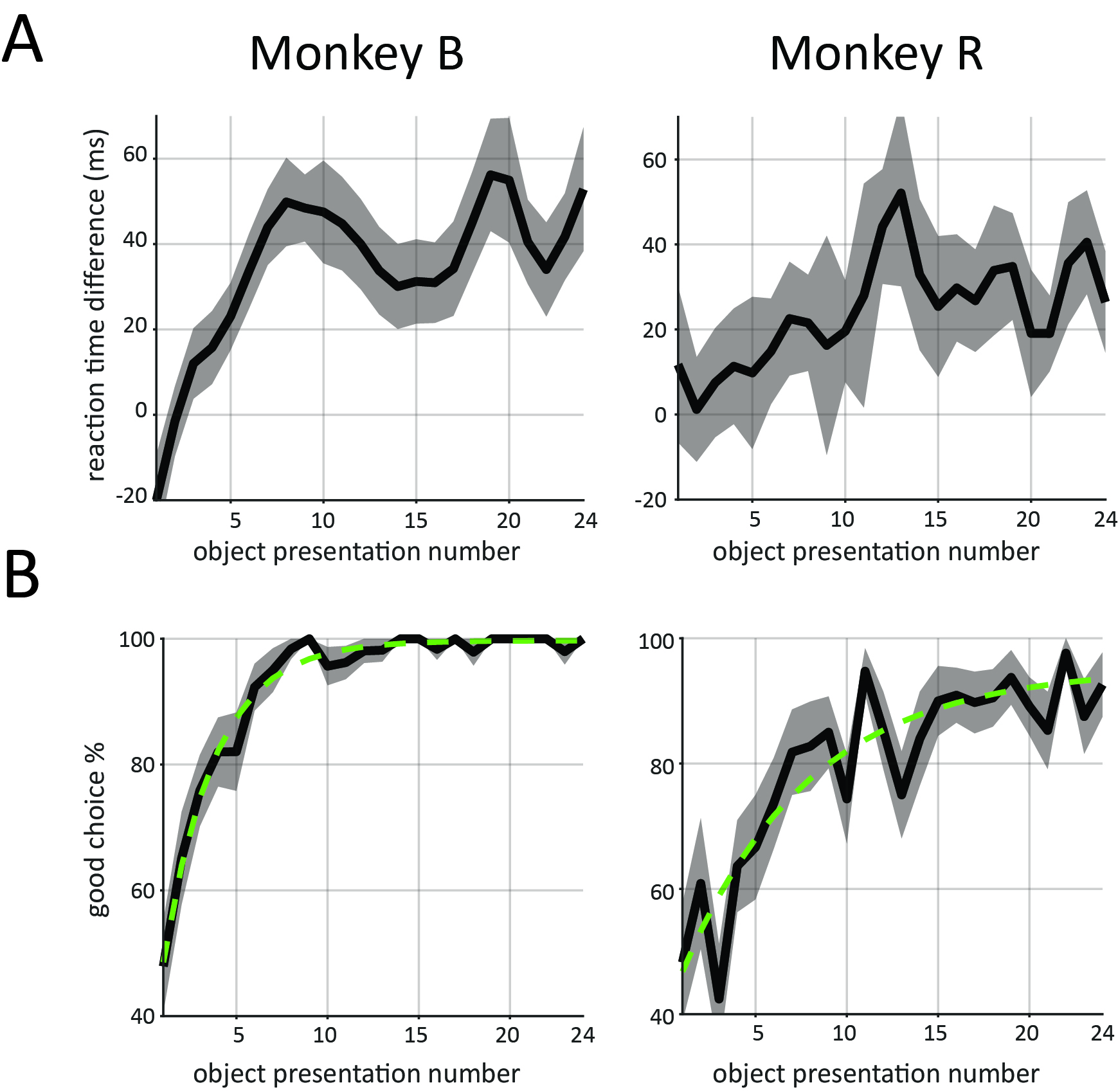

### Supplemental Figure 2

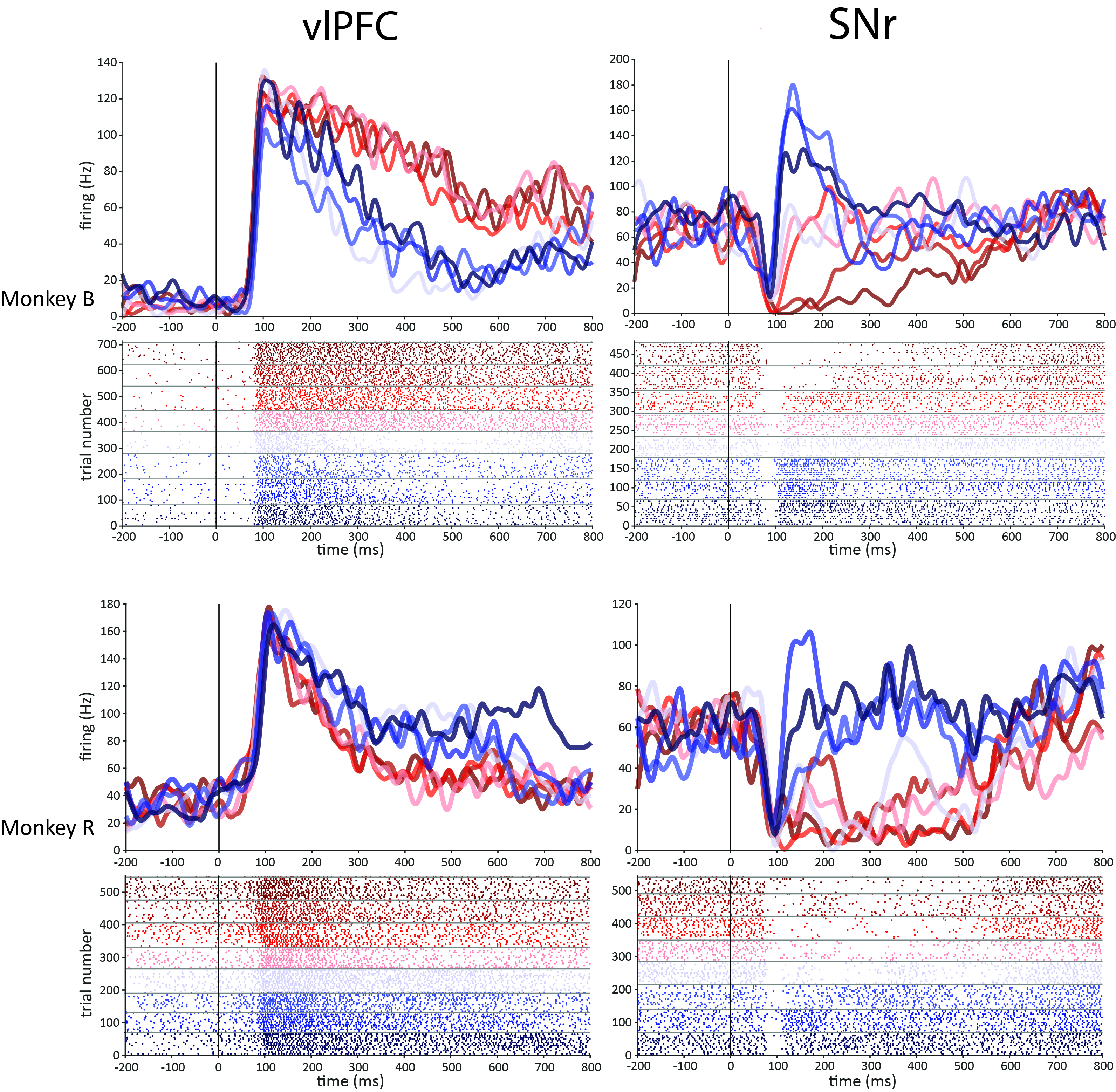

### Supplemental Figure 3

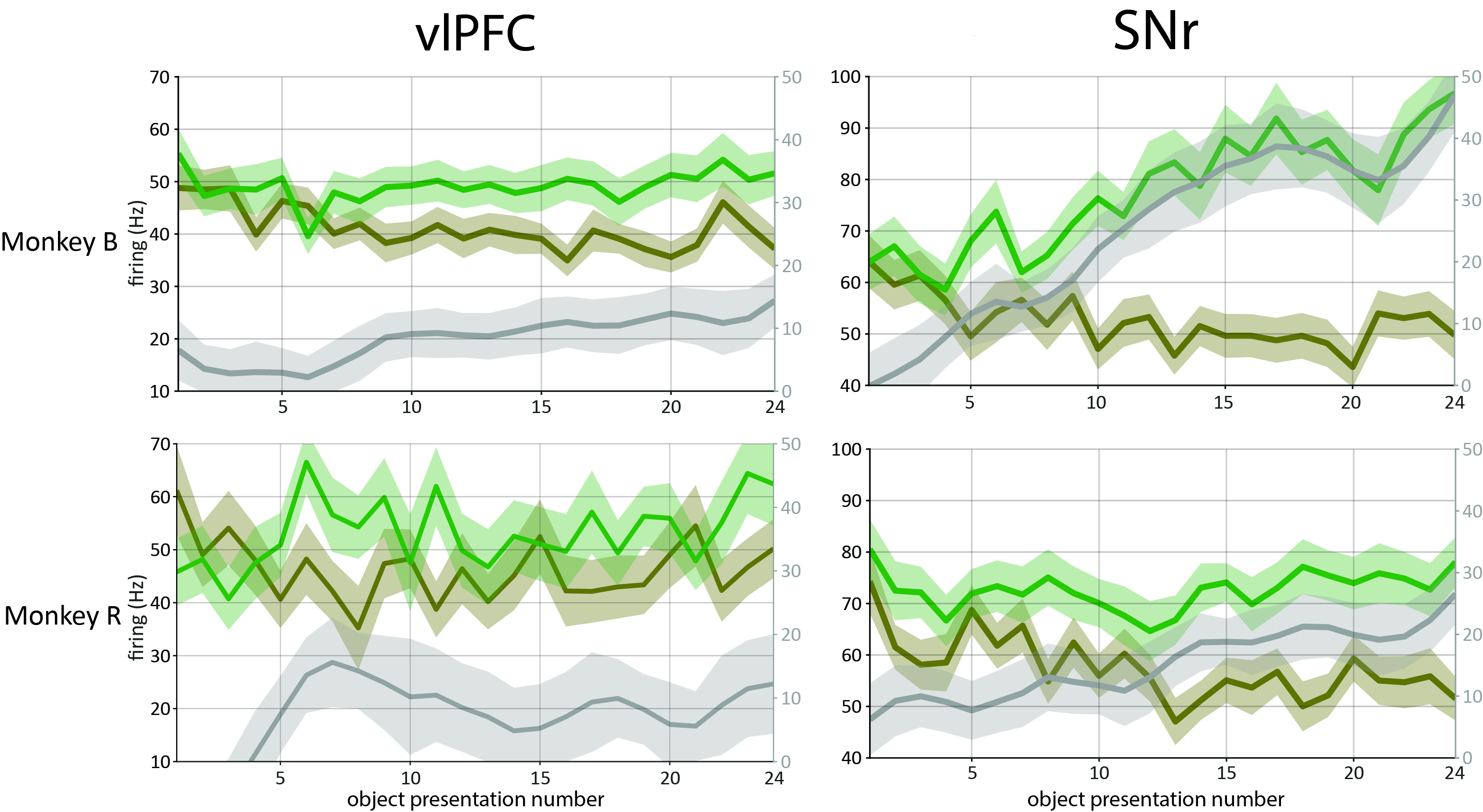

### Supplemental Figure 4

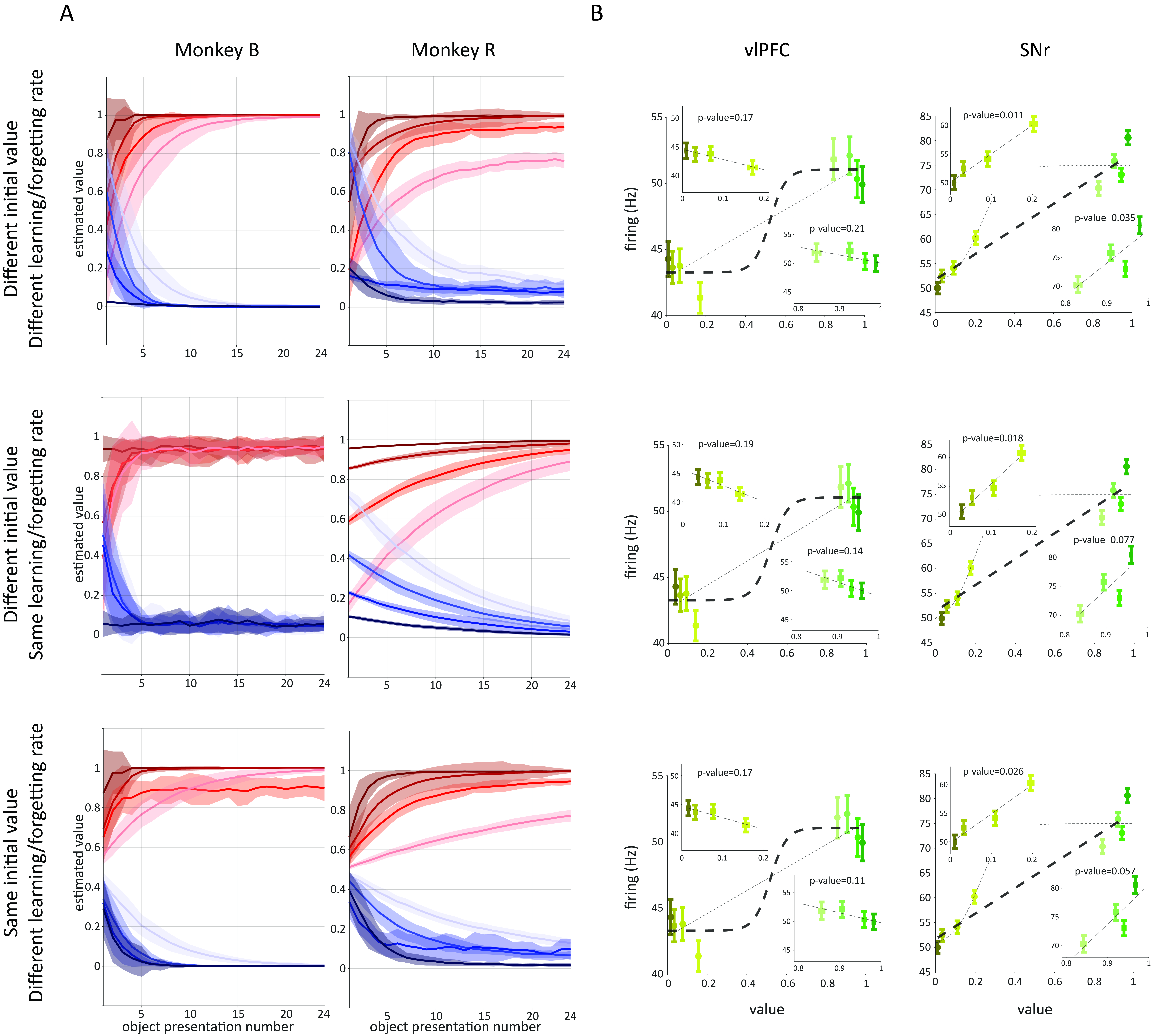

### Supplemental Figure 5

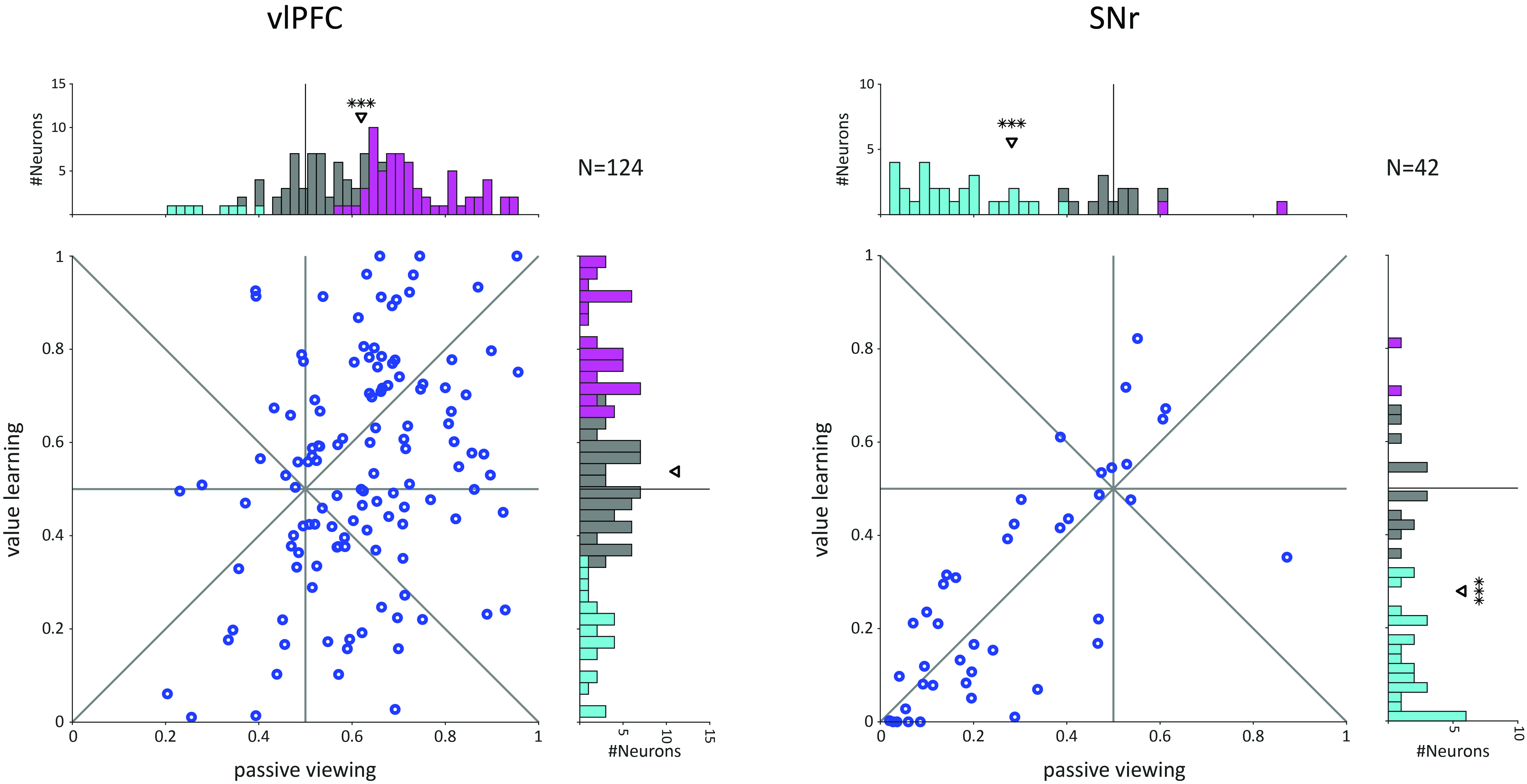
